# Are we missing the forest for the trees? Conspecific negative density dependence in a temperate deciduous forest

**DOI:** 10.1101/2021.01.06.425540

**Authors:** Kathryn E. Barry, Stefan A. Schnitzer

## Abstract

One of the central goals of ecology is to determine the mechanisms that enable coexistence among species. Evidence is accruing that conspecific negative density dependence (CNDD), the process by which plant seedlings are unable to survive in the area surrounding adults of their same species, is a major contributor to tree species coexistence. However, for CNDD to maintain diversity, three conditions must be met. First, CNDD must maintain diversity for the majority of the woody plant community (rather than merely specific groups). Second, the pattern of repelled recruitment must increase in with plant size. Third, CNDD must occurs across life history strategies and not be restricted to a single life history strategy. These three conditions are rarely tested simultaneously. In this study, we simultaneously test all three conditions in a woody plant community in a North American temperate forest. We examined whether the different woody plant growth forms (shrubs, understory trees, mid-story trees, canopy trees, and lianas) at different ontogenetic stages (seedling, sapling, and adult) were overdispersed – a spatial pattern indicative of CNDD – using spatial point pattern analysis across life history stages and strategies. We found that there was a strong signal of overdispersal at the community level. However, this pattern was driven by adult canopy trees. By contrast, understory plants, which can constitute up to 80% of temperate forest plant diversity, were not overdispersed as adults. The lack of overdispersal suggests that CNDD is unlikely to be a major mechanism maintaining understory plant diversity. The focus on trees for the vast majority of CNDD studies may have biased the perception of the prevalence of CNDD as a dominant mechanism that maintains community-level diversity when, according to our data, CNDD may be restricted largely to trees.

## Introduction

Conspecific negative density dependence (CNDD) is one of the most empirically supported mechanisms for the maintenance of plant species diversity (Mangan *et al*. 2010, Johnson *et al*. 2012; Comita *et al*. 2014, La Manna *et al*. 2017). Conspecific negative density dependence occurs when small individuals have relatively low rates of growth and survival near adult members of their own species (conspecifics). This constraint on growth near members of the same species results in a distinct spatial pattern where adult conspecifics occur further away from each other than would be expected by chance (overdispersion; Ledo & Schnitzer 2014; but see Murrell 2009, LaManna et al. 2017). Thus, CNDD is predicted to result in stable species coexistence across the landscape because dominant species cannot displace subordinate ones (Janzen 1970; Connell 1971). Evidence for CNDD has been reported in a variety of ecosystems, including lakes, deserts, grasslands, marine ecosystems, and particularly in temperate and tropical forests (Anderson 2001; Goldberg *et al*. 2001; Lorenzen & Enberg 2002; Petermann *et al*. 2008; Comita *et al*. 2010; Mangan *et al*. 2010; Schnitzer *et al*. 2011; Johnson *et al*. 2012; Johnson *et al*. 2014, LaManna et al. 2017). For forests, over the past ten years alone, evidence for CNDD has been reported more than 30 times in 13 countries across five continents (Barry & Schnitzer 2016).

However, strong evidence for CNDD at the seedling level may not maintain diversity at the forest level if NDD reduces clustering of seedlings but does not overcome the initial clumped pattern of seedlings around adults (due to dispersal limitation) (Zhu *et al*. 2015). That is, if CNDD does not result in a pattern of overdispersion, then it may fail to stably maintain species diversity (Hubbell 1979; 1980; Ledo & Schnitzer 2014; but see Muller-Landau & Adler 2007). For CNDD to maintain community-level diversity in temperate forests, the following three conditions must be met. 1) Individuals of the majority of the species in the community will be overdispersed because of greater mortality near conspecific adults. If only a few species are overdispersed, then CNDD will not theoretically create a sufficient rare species effect to maintain diversity (Janzen 1970, Connell 1971). 2) The degree of overdispersion will increase with life-history stage. That is, the signal of CNDD should compound as individuals mature, and thus larger individuals of any given species should be more overdispersed than smaller individuals (see Zhu *et al*. 2015). 3) CNDD will operate across life history strategies, including growth form and dispersal syndrome. In particular, CNDD will operate in the plant growth forms that have the highest diversity if it is the main mechanism driving diversity maintenance, as suggested by previous studies (reviewed by Comita *et al*. 2014, Barry & Schnitzer 2016). If these three conditions are met, then CNDD is sufficiently strong to decrease initial clustered dispersal patterns and to maintain community-level diversity.

While there is abundant evidence for CNDD as a diversity maintenance mechanism in vascular plant communities, to date, no single study has tested the three necessary conditions to confirm that NDD maintains plant species diversity in any ecosystem. Previous studies may have overestimated the importance of CNDD in forest ecosystems for two reasons. First, the vast majority of studies that examined CNDD in vascular plant species focused on growth and mortality at the seedling stage (Packer & Clay 2000; 2003; Comita *et al*. 2010; Mangan *et al*. 2010; Johnson *et al*. 2012; Johnson *et al*. 2014). Dynamics at the seed-to-seedling and seedling-to-sapling transitions do not necessarily translate to overdispersion in the larger size classes (Yao et al. 2020) and may overestimate CNDD (Detto et al. 2019). That is, dispersal limitation results in most seeds arriving underneath the parent tree and CNDD dynamics at the seed-to-seedling, and seedling-to-sapling transitions must be sufficiently strong to overcome the spatial signature of dispersal limitation. The negative effects of growing near a conspecific adult should compound as individuals mature, and thus, the level of overdispersion should increase with plant size. Second, the vast majority of CNDD studies in forests focused only on canopy trees, ignoring other important plant growth forms (e.g., Ledo & Schnitzer 2014). The biased selection in growth form is particularly problematic in temperate forests, where canopy tree species represent a relatively small fraction (∼20%) of the total vascular plant community (Gilliam 2007). Furthermore, canopy trees may be more prone to overdispersion due to their capacity for long distance dispersal (McCarthy-Neumann & Kobe 2008; Kobe & Vriesendorp 2011; DeWalt *et al*. 2015). Our perception of the prevalence of CNDD and overdispersion may be biased by the focus on canopy trees. By contrast, understory plants have a lower capacity for long distance dispersal due to their relatively short stature and position in intact forests, and thus may be less likely to be overdispersed. Nonetheless, if CNDD is the primary mechanism that maintains diversity, we would expect it to operate across life history stage and life history strategy.

We addressed each of the three core conditions for CNDD to be a general mechanism for the maintenance of plant species diversity by evaluating the spatial patterns of a woody plant community across life history strategies (shrubs, understory trees, mid-story trees, canopy trees, and lianas) and stages (seedling, sapling, and adult) in a temperate forest in western Pennsylvania, USA. We tested three specific hypotheses: 1) Woody plant diversity in temperate forests is maintained by CNDD, and thus we predict that the majority of plant species will be overdispersed. 2) The effects of CNDD compound as plants mature (across ontogeny), and thus we predict that overdispersion will increase with plant size. 3) CNDD operates independently of growth form and life history strategy, and thus we predict that the pattern of overdispersion will be found in the majority of the species of all plant groups. We tested these hypotheses by examining the degree of overdispersion in a woody plant community, which included a range of plant life-history stages (*i*.*e*., sizes) and life-history strategies (*i*.*e*., growth form and dispersal syndrome).

## Materials and Methods

### Study site

We conducted this study at Powdermill Nature Reserve, an 890-hectare reserve located in the Allegheny plateau at the base of the Appalachian Mountains in southwestern Pennsylvania, USA (Westmoreland County; 40°09’ N, 79°16’ W). This region receives ∼1100 mm of precipitation per year and is characterized by mixed mesophytic vegetation that is dominated by maples (*Acer spp*.), tuliptree (*Liriodendron tulipifera*), and oaks (*Quercus spp*., Murphy *et al*. 2015). Elevation at Powdermill Nature Reserve ranges from 392 to 647 m above sea level. Powdermill Nature Reserve contains a matrix of vegetation types consisting primarily of secondary deciduous forest but with several areas of maintained fields and managed lands. Last known logging occurred in this region in the 19th century, and land was primarily used for agriculture into the early 20th century (see Murphy *et al*. 2015 for more detailed site description).

### Plot establishment and plant census

We established sixteen 10-m diameter circular plots in the >90-year-old secondary temperate deciduous forests at Powdermill Nature Reserve. We chose the 10-m diameter spatial grain because this size was thought to be a suitable size to test for spatial patterns associated with NDD in a Malaysian forest (Zhu *et al*. 2013; see also Hubbell 1979, Condit *et al*. 1992). We avoided canopy gaps for the placement of each plot, and each plot had >80% canopy cover. We ensured that the plots were not within 10 m of a waterway, that soil cover was not predominantly rocks, and that plots were at least 20 m from any edge. We used a Trimble GeoExplorer 6000XH to measure the precise location (up to 10 cm accuracy) of all woody plant individuals >10 cm height in each plot (Trimble Navigation Limited, Westminster, CO). For each individual, we measured height and basal diameter, and we identified them to species.

To examine how overdispersion changes with plant size, we divided individuals into four height classes (<0.5 m, 0.5-1 m, 1-5 m, and 5-10 m). To understand how overdispersion interacts with life-history strategy, we classified each species as either canopy or understory, and as either bird, wind, self, or other animal dispersed based on species descriptions in the Flora of North America (Flora of North America Editorial Committee eds. 1993).

### Data analysis

We performed all data analysis in R statistical computing software (v. 3.2.2, R Development Core Team 2015). To measure plant spatial distribution (the degree to which plants are clustered or overdispersed), we calculated Ripley’s K in the package “spatstat” using Ripley’s translational border correction at each plot for each species and then converted K to Besag’s L (Ripley 1977; Besag 1977; Baddeley & Turner 2005; Baddeley *et al*. 2015). Several studies have demonstrated that spatial point pattern analysis is capable of detecting spatial patterns that can be attributed to mechanistic processes (e.g. Ledo & Schnitzer 2014; Brown *et al*. 2015). To eliminate point patterns based on low replication, we removed species at any plot with fewer than five individuals (resulting in final data from a total of 16 plots). We then calculated a pooled L for each comparison (by species, by species type, by species type/plant size, or by species type/dispersal mechanism) by weighting the individual L estimates by the number of points in a given L-function (methods follow Bagchi & Illian 2015). We bootstrapped these estimates 999 times to create 95% confidence intervals. We then calculated the predicted L for complete spatial random to ensure that reported spatial patterns were significantly different from (not overlapping with) complete spatial random. Data manipulation of input to and output from point pattern analysis was done using a combination of the “abind”, “gridExtra”, and “reshape” packages (Wickham 2007; Aguie 2015; Plate & Heiberger 2015). We constructed all figures in the package “ggplot2” (Wickham 2009).

To more easily interpret the figures, we corrected our measures of L with the distance at which each measure of L is calculated (L(d)-d). Besag’s L is a measure of spatial aggregation, and when L(d)-d is positive, a greater proportion of neighbors are observed within distance d of focal individuals than predicted by a complete spatial random pattern. When L(d)-d is negative, a smaller proportion of neighbors are observed within distance d of focal individuals than predicted by a complete spatial random pattern (Besag 1977). We classify any point pattern where a linear regression slope of L(d)-d is significant and positive (using the command “lm” in R, Table S1, S2) with increasing distance (d) as overdispersed (Bagchi & Illian 2015). This designation implies that more individuals are found far away from an individual of a given species than near an individual of that species. We classify any point pattern where a linear regression slope of L(d)-d is significant and negative with increasing distance as clustered. These designations differ from “pure overdispersion” (i.e. regularity or inhibition), which would begin with a significantly negative L(d)-d that indicates fewer individuals close to the parent than would be expected by chance (Dale 1999; Bagchi & Illian 2015; Baddeley *et al*. 2015). However, natural dispersal typically results in more conspecific seeds and seedlings close to adults than predicted by complete spatial random, and thus we did not expect to find a significant negative L(d)-d of seedlings close to the parent (van der Pijl 1982). Therefore, we accounted for dispersal limitation by focusing on overdispersion as having a positive slope with regards to distance (d), which indicates a significant increase in individuals with distance from the adult, at the community scale. However, CNDD should result in increasing overdispersion with plant size as the effects of CNDD compound with time (as the plant matures). Thus, we might expect a spatial signature of “pure overdispersion” in larger size classes, if CNDD is capable of overcoming initial dispersal patterns and thus stably maintaining coexistence.

We considered any point pattern to be significantly different from complete spatial random if a mixed effect linear model (calculated using the command “lmer” in package “lme4” with plot as a random effect and using the package “lmerTest” to calculate p-values) of the L(d)-d and the distance was significant (Table 1; Bates *et al*. 2015, Kuznetsova *et al*. 2015). If the total model was considered significant, we did not consider the point pattern to be significantly different from complete spatial random at any distance where the bootstrapped 95% confidence intervals of L(d)-d overlap with complete spatial random. We considered any two point patterns to be significantly different from each other if their bootstrapped 95% confidence intervals did not overlap at a given distance.

**Table 1:** **Results of mixed effects linear model to calculate L statistic significance at Powdermill Nature Reserve in Southwestern Pennsylvania for all comparisons of all individuals >10 cm height**. To calculate significant differences from complete spatial random we used a mixed effects linear model with plot as a random effect to control for between plot differences due to environmental heterogeneity between plots. We report model degrees of freedom based on the number of L estimates (calculated every 10 cm per point pattern per plot).

| **Model** | **dF** | **T Stat** | **P value** | **Figure** |
| --- | --- | --- | --- | --- |
| All individuals | 2228 | 19.37 | <0.0001 | 1a |
| All individuals, <0.5 | 781 | -4.477 | 0.6689 | 1b |
| All individuals, 0.5-1 | 889 | 7.83 | <0.0001 | 1b |
| All individuals, 1-5 | 797 | 8.5828 | <0.0001 | 1b |
| All individuals, >5 | 358 | -14.97 | <0.0001 | 1b |
| Understory | 1071 | 10.191 | <0.0001 | 2a |
| Canopy | 1059 | 5.796 | <0.0001 | 2a |
| Canopy, <0.5 m tall | 380 | -7.14 | <0.0001 | 2b |
| Canopy, 0.5-1 m tall | 478 | 8.67 | <0.0001 | 2b |
| Canopy, 1-5 m tall | 463 | 9.14 | <0.0001 | 2b |
| Canopy, >5 m tall | 186 | -10.71 | <0.0001 | 2b |
| Understory, <0.5 m tall | 397 | 2.045 | 0.0415 | 2c |
| Understory, 0.5-1 m tall | 407 | 0.3348 | 0.7379 | 2c |
| Understory, 1-5 m tall | 381 | 1.038 | 0.2998 | 2c |
| All individuals, bird dispersed | 1159 | 2.86 | 0.0042 | 3a |
| All individuals, animal dispersed | 200 | -6.806 | <0.0001 | 3a |
| All individuals, wind dispersed | 803 | 3.404 | <0.0001 | 3a |
| Overstory, wind dispersed | 601 | 3.903 | <0.0001 | 3b |
| Overstory, bird dispersed | 250 | 5.482 | <0.0001 | 3b |
| Understory, wind dispersed | 199 | -8.895 | <0.0001 | 3b |
| Understory, bird dispersed | 906 | -3.533 | 0.0004 | 3b |

## Results

At the community level, all woody plants combined were significantly overdispersed (Figure 1a). The largest individuals (>5 m tall) had a significantly lower dispersion (L(d)-d) at intermediate distances (2-5m), than the two middle height size classes (1m – 5m and 0.5 – 1 m); however, L(d) – d did not differ significantly among the larger size classes at distances greater than 5 m (Figure 1b). By contrast, the smallest individuals (< 0.5 m) had significantly lower overdispersion than intermediate height individuals (0.5-1m and 1-5 m tall) for all distances greater than 2m (Figure 1b), and significantly lower dispersion than individuals in all of the larger height categories for distances greater than 5m. Thus, all but the smallest size classes were overdispersed at longer distances from the adult tree, indicating that, at the community-level, NDD was strong enough to overcome the initial clumped distribution of seedlings.

**Figure 1.**
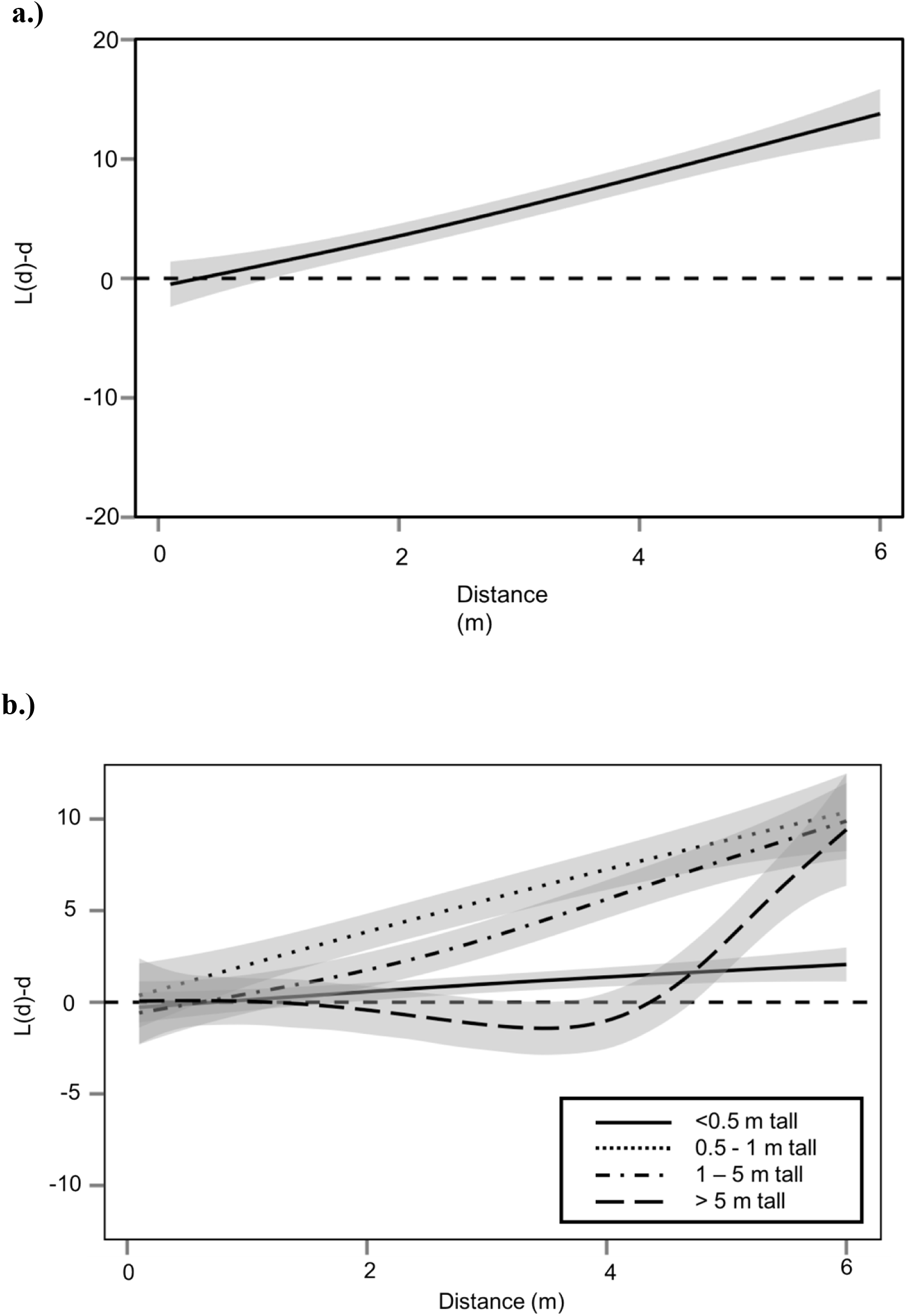
Pooled Besag’s L statistic across distance from spatial point pattern analysis for the full community of woody plants >10 cm in height at Powdermill Nature Reserve in Southwestern Pennsylvania. a.) The community of woody plants (all species, n=62 point patterns) was significant overdispersed regardless of dispersal mechanism. However, the L(d)-d for the community remains positive across all distances indicating that some individuals occur close to members of their own species. b.) Individuals that were <0.5 m tall were the least overdispersed (n=25 point patterns). Individuals that were intermediate in height (0.5m to 5 m tall) were significantly more overdispersed than smaller individuals, though not significantly more or less overdispersed than the largest individuals (n_0.5-1m_=26 point patterns, n_1-5m_=27 point patterns). The largest individuals (> 5m tall, n= 13 point patterns) were not significantly more overdispersed than individuals that were 0.5m to 5m tall; however, the drop in the line below complete spatial random indicates that they had less clumping over small distances. Grey shaded regions represent 95% confidence intervals, darker grey regions represent overlapping confidence intervals.

Both canopy trees and understory plants were significantly overdispersed; canopy trees were more overdispersed (significantly higher L(d)-d) at distances greater than 3 m (Fig. 2a). The differences in overdispersion between canopy and understory plants become more pronounced with plant life history stage (*i*.*e*., plant size). Canopy trees did not differ significantly from complete spatial random when they were small and young, but became significantly overdispersed when they were larger (Figure 2b), which is consistent with CNDD. Understory plants displayed the opposite pattern: they were overdispersed when small, but larger individuals were indistinguishable from complete spatial random (Figure 2c).

**Figure 2:**
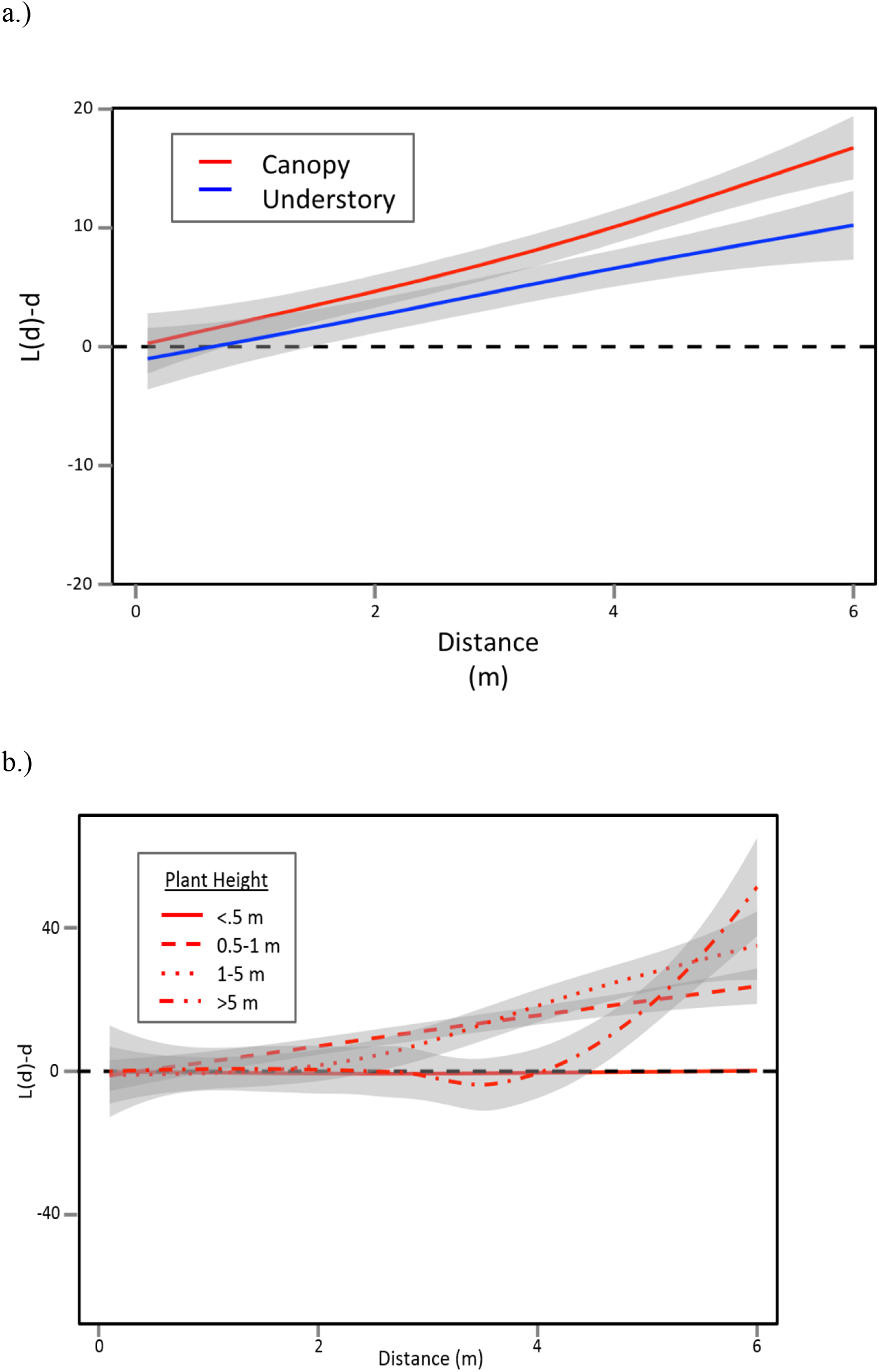
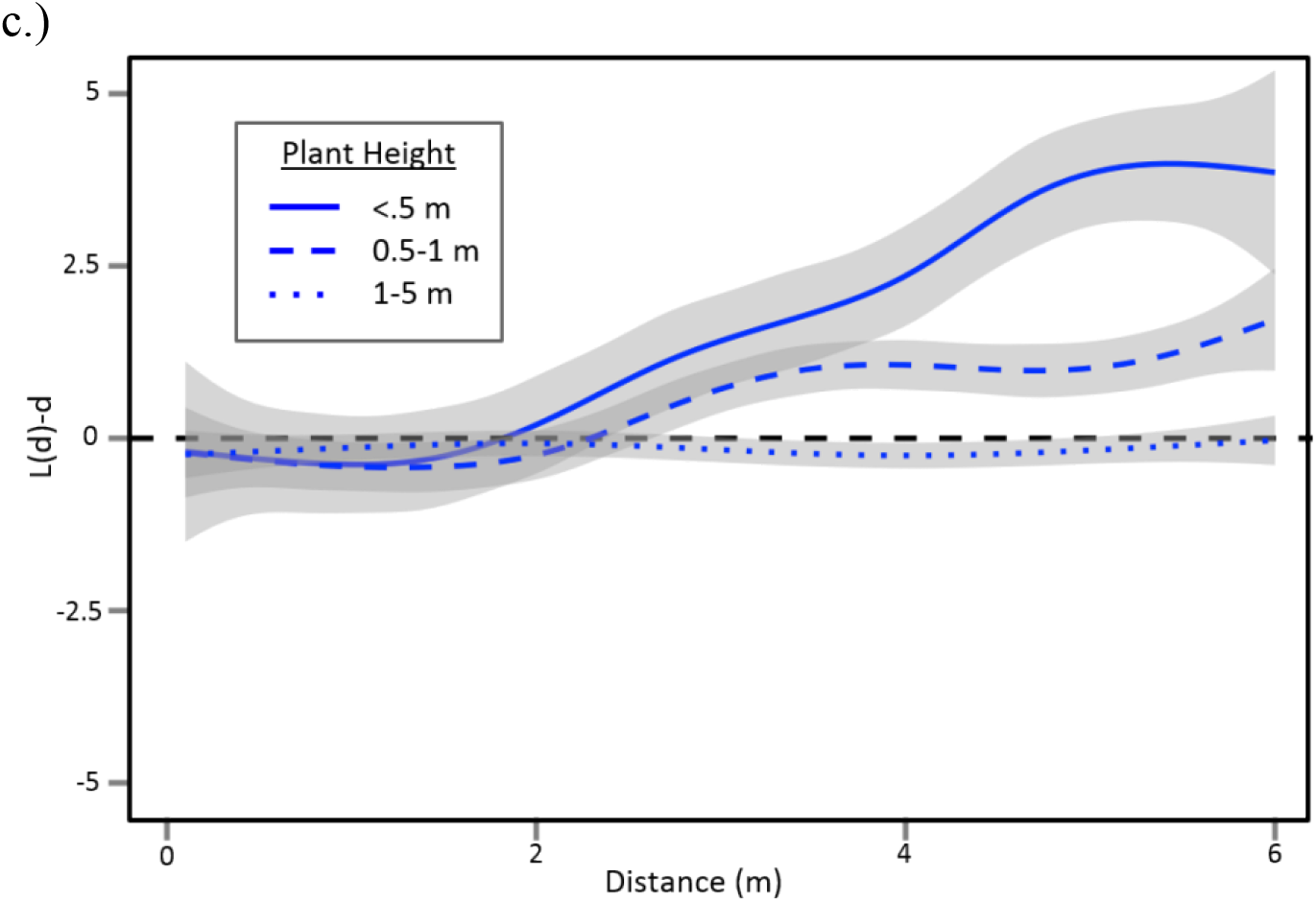
Pooled Besag’s L statistic across distance from spatial point pattern analysis for woody plants >10 cm in height at Powdermill Nature Reserve in Southwestern Pennsylvania separated by growth form. a.) Canopy(n = 29 point patterns) and understory plants (n=33 point patterns) were both significantly overdispersed, indicative of negative density dependence. Canopy plants were significantly more overdispersed than understory plants. b.) Canopy plants were more overdispersed with increasing life-history stage in accordance with predictions for negative density dependence (n_<0.5_=14 point patterns, n_0.5-1_=48 point patterns, n_1-5_=32 point patterns, n_>5_=15 point patterns). c.) Understory plants were not more overdispersed with life-history stage (n_<0.5_=21 point patterns, n_0.5-1_=22 point patterns, n_1-5_=20 point patterns). Grey shaded regions represent 95% confidence intervals, darker grey regions represent overlapping confidence intervals.

All of the four dispersal mechanisms that we examined, wind, bird, and self-dispersed species were overdispersed and statistically indistinguishable from each other. Species dispersed by animals other than birds (including secondary dispersal by squirrels) were all significantly less overdispersed than the other three dispersal types (Figure 3a). Dispersal syndrome for bird and wind dispersed species did not explain the differences in spatial pattern between canopy trees and understory plants; canopy trees were always more overdispersed than understory plants regardless of dispersal mechanism (Figure 3b), suggesting that the height of canopy trees is the most important factor in dispersal distance. We limited this analysis to bird and wind dispersed species because these two groups had sufficient replication for robust comparisons between canopy and understory plants.

**Figure 3.**
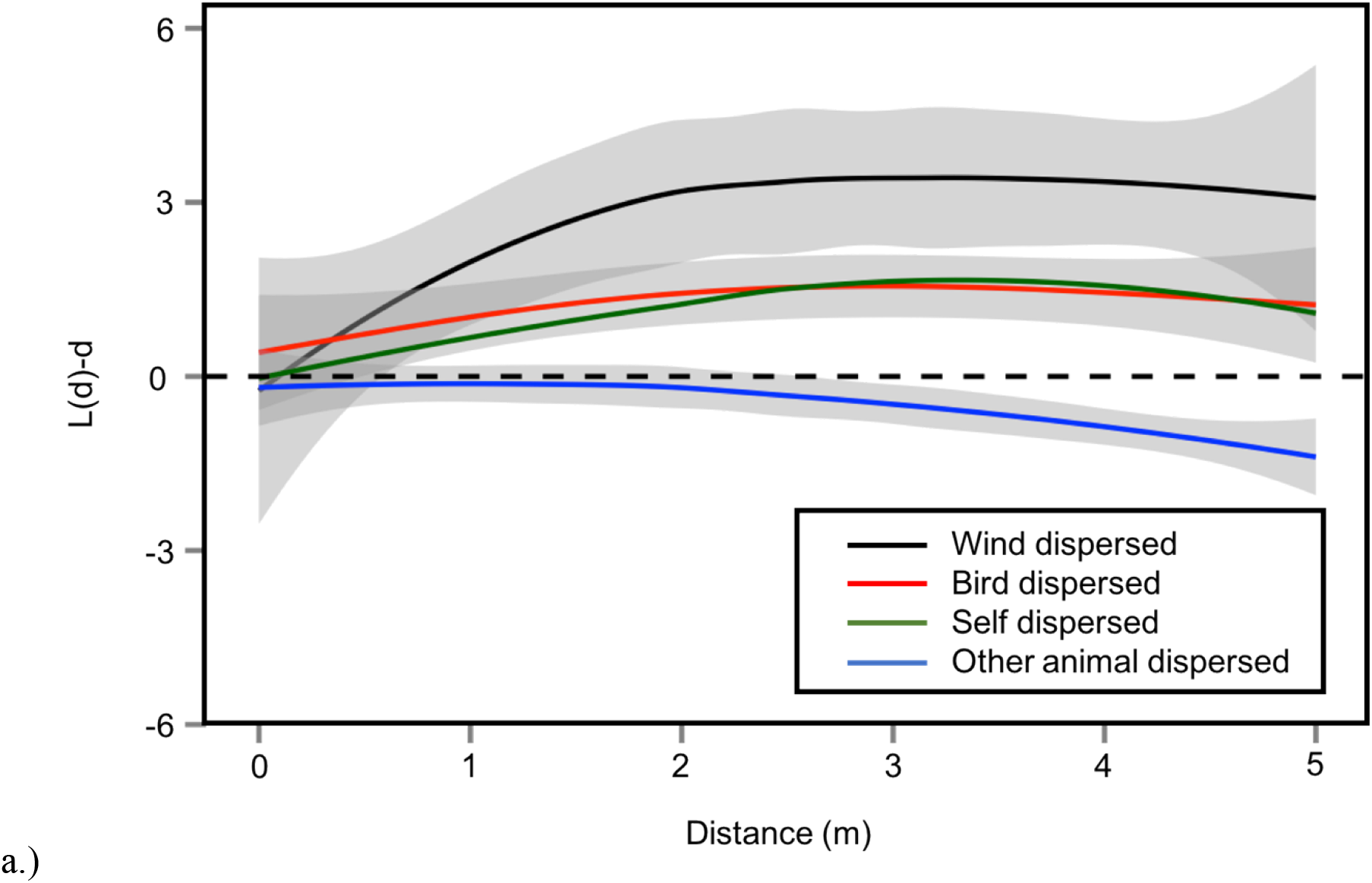
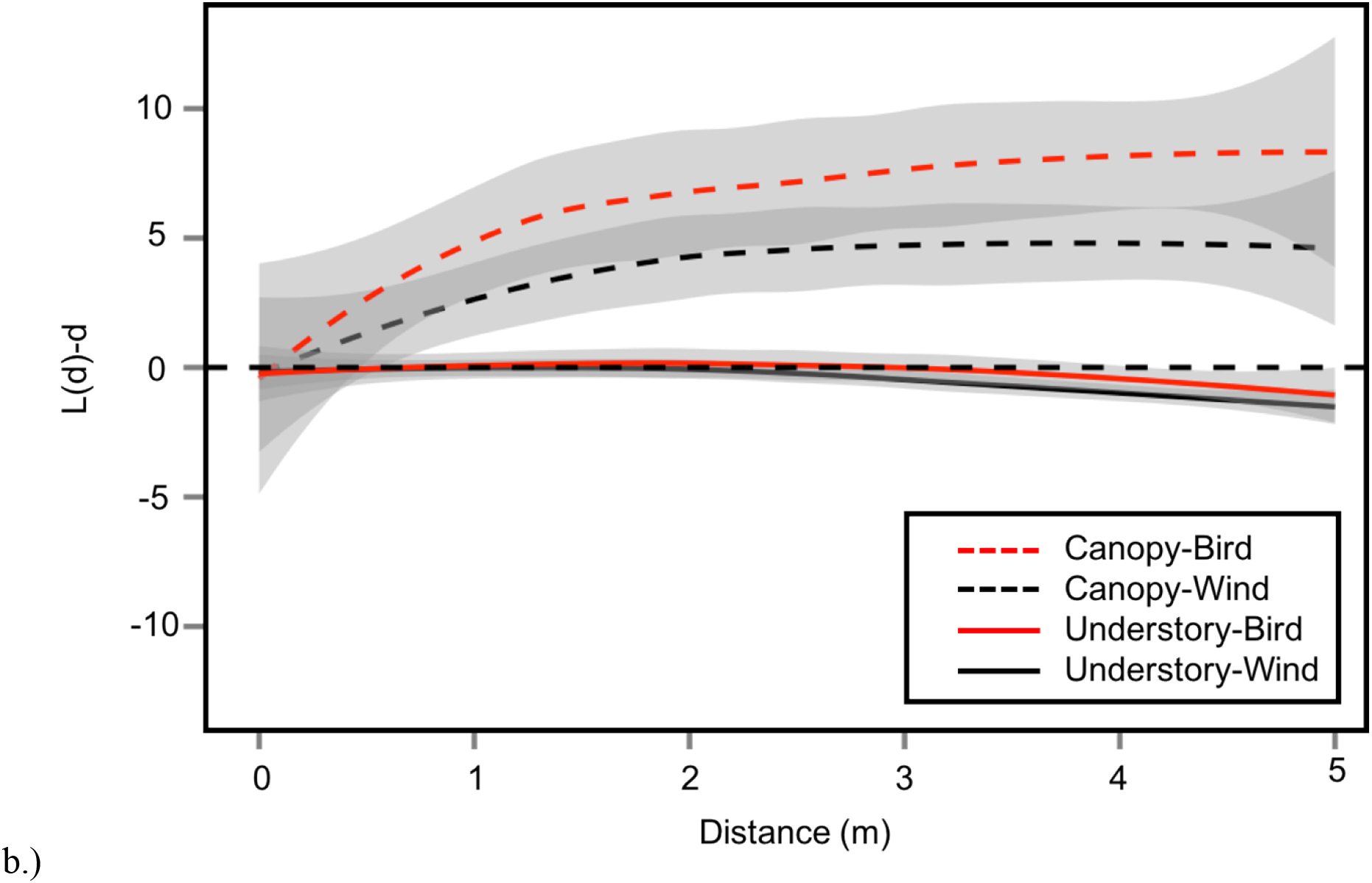
Pooled Besag’s L statistic across distance from spatial point pattern analysis of the woody plant community stratified by dispersal mechanism and plant type at Powdermill Nature Reserve in southwestern Pennsylvania. a.) Wind dispersed(n=16), bird dispersed (n=23), and self dispersed(n=2) species were significantly more overdispersed than species dispersed by animals other than birds(n_animal_=6). b.) Canopy plants were significantly more overdispersed than understory plants regardless of dispersal mechanism(n_canopy-bird_=5, n_canopy-wind_=12, n_understory-bird_=18, n_understory-wind_=5). Bird dispersal was emphasized here; however, plants did not differ significantly from wind dispersed plants either in the canopy and the understory. All reported sample sizes (n) are in number of total point patterns contributing to a pooled L function. Grey shaded regions represent 95% confidence intervals, darker grey regions represent overlapping confidence intervals.

## Discussion

We found that canopy trees were overdispersed and the strength of overdispersion increased with tree size - two critical conditions for CNDD to be a mechanism that maintains species diversity. These findings are consistent with a growing number of studies that have reported that CNDD is a viable mechanism to maintain canopy tree diversity in temperate and tropical forests (*e*.*g*., Comita *et al*. 2010; Mangan *et al*. 2010; Johnson *et al*. 2012; Zhu *et al*. 2015; reviewed by Comita *et al*. 2014, LaManna et al. 2016, LaManna et al. 2017). Thus, our findings support CNDD as a mechanism for the maintenance of canopy tree species diversity.

For woody understory plants, however, our spatial patterns did not meet the criteria for CNDD to maintain species diversity at Powdermill Nature Reserve. Understory species were overdispersed only in the smallest size classes, and overdispersion did not increase with plant life-history stage. For CNDD to be a viable diversity maintenance mechanism, overdispersion should increase with plant size, as we found for canopy trees. Thus, had CNDD been present in understory plants, it was not sufficiently strong to overcome the initial clumped dispersal pattern of seedlings, and therefore it did not result in overdispersion. The lack of sufficient strength to overcome dispersal patterns indicates that CNDD does not appear to produce stable coexistence in these species. Similar conclusions that CNDD is not a general mechanism for the maintenance of non-tree plant diversity were reported for tropical forests. For example, Ledo & Schnitzer (2014) found that lianas, which comprised ∼35% of the woody species diversity in a Panamanian tropical forest (Schnitzer *et al*. 2012, 2015), were underdispersed (clustered) rather than overdispersed. Thus, Ledo & Schnitzer (2014) concluded that, while there was evidence for CNDD for canopy trees, there was little evidence for CNDD for lianas, indicating that CNDD was relevant for trees, but was not a general mechanism for the maintenance of woody plant species diversity in their forests. In a Caribbean tropical forest, DeWalt and colleagues (2015) found that non-canopy tree woody seedlings (lianas and shrubs) were less likely to suffer negative density dependent mortality than canopy trees. In tropical forests, however, trees commonly represent 65% or more of the woody plant species diversity (*e*.*g*., Schnitzer *et al*. 2012; 2015), and thus CNDD is still likely a powerful diversity maintenance mechanism. By contrast, CNDD may fail to maintain the majority of species diversity in temperate forests where understory species can represent the majority (∼80%) of the diversity (Gilliam 2014).

It is likely that several mechanisms are acting in concert to maintain community-level diversity. This combination of mechanisms may occur broadly across plant groups as suggested by Ledo and Schnitzer (2014), where the authors found that clumped spatial distributions may be due to niche regeneration in lianas, which differs from the mechanism(s) maintaining the majority of tropical trees. Similarly, Yao et al. (2020) found that CNDD was likely important for individuals when they were young and small but that topographic and edaphic factors increased in importance with increasing plant age.

In our temperate forest, it seems likely that several mechanisms are occurring simultaneously for canopy and understory plants. Trees may be maintained largely by CNDD; whereas, understory plants may be influenced by a number of different mechanisms. There is evidence that CNDD is a weak mechanism for the maintenance of understory plant diversity, since overdispersion is present when understory plants are small (Fig. 2c). However, the lack of overdispersion in larger understory plants indicates that a mechanism (or mechanisms) other than CNDD is a stronger driver of understory plant diversity. These factors may include abiotic factors as found by both Ledo and Schnitzer (2014) and Yao et al. (2020).

Canopy trees may be significantly more overdispersed than understory species simply because being tall enables longer distance dispersal. We found higher overdispersion of canopy trees than for understory plants regardless of dispersal mechanism (Figure 3b). Plant height rather than dispersal mechanism appeared to account for greater overdispersion in canopy trees. Understory plants tend to have universally smaller dispersal kernels regardless of dispersal mechanism because of their smaller stature (van der Pijl 1982). Small stature results in fewer seeds dispersed at longer distances - even for bird-dispersed seeds (Figure 3b). The inability to move seeds far away from the parent tree may force understory plants to be better defended against soil pathogens, which appear to be strong agents of CNDD (Bever 2003; Packer & Clay 2003; Kulmatiski *et al*. 2008; Mangan *et al*. 2010). Furthermore, negative feedback from soil pathogens may be inversely related to light availability (Smith & Reynolds 2015, Jiang et al. 2020). Indeed, many understory plants are naturally well defended because of the importance of preserving plant tissue in a low-light environment (Coley 1983; Coley *et al*. 1985); thus, understory plants may be predisposed to developing greater defenses to pathogens rather than increasing dispersal abilities.

Differences in the level of overdispersion between canopy species and understory species did not appear to be due to the spatial scale of study in spite of our relatively small plot size. If spatial scale had biased our results, we would have expected the spatial point pattern analysis to show little evidence of overdispersion for large canopy trees, but rather a signature indistinguishable from complete spatial random. Furthermore, Zhu *et al*. (2010) demonstrated that when NDD is present it is most likely to be present at the 0-5 m scale and peaks at 5 m. Our results showed a clear spatial signature of overdispersion for our largest individuals. Thus, it seems unlikely that our findings were caused by differences in plant scale. Furthermore, Bagchi & Illian (2015) demonstrate that replicated point pattern analysis is significantly more robust to problems of small scale than traditional point pattern analysis.

## Conclusions

The intense focus on canopy trees, and in particular on tree seedlings, may bias the current understanding of diversity maintenance in forest ecosystems (Detto et al. 2019, Yao et al. 2020). If we had restricted our sampling to only the smallest understory individuals, we would have concluded that CNDD maintains woody understory plant diversity but not canopy tree diversity. The smallest understory individuals (<0.5 m tall) were overdispersed, while the smallest canopy individuals did not differ significantly from complete spatial random (Figure 2). However, examining larger individuals indicated that adult canopy trees became overdispersed as they matured, but that understory plants did not. Zhu and colleagues (2015), Detto and colleagues (2019) and Yao and colleagues (2020) all emphasized similar caution in drawing large-scale conclusions from studies of seedling dynamics for three reasons. First, patterns of seedling mortality often have little effect on broader community and demographic patterns (Zhu et al. 2015). Second, NDD tends to decrease with ontogeny rather than increase (Yao et al. 2019). Finally, studies of NDD at the recruitment level may overestimate NDD due to regression dilution (Detto et al. 2019).

To fully understand the maintenance of plant species diversity, it is necessary to examine spatial patterns across plant sizes, as well as across plant groups that vary in life history strategies. Spatial patterns may be even more complex when considering species that vary more broadly in their life history strategies, such as herbaceous species, which comprise the majority of plant diversity in temperate deciduous forests (Gilliam 2007) and are largely neglected with regard to their diversity maintenance (Barry & Schnitzer 2016). Nevertheless, even by simply dividing the woody plant community into canopy trees and woody understory plants, we demonstrate that CNDD, which appears to maintain canopy tree diversity, may not be strong enough to overcome dispersal limitation and maintain understory woody plant diversity in this temperate forest.

## Author contributions

K.E.B provided initial project idea, collected and analyzed data, and wrote initial drafts of this paper. S.A.S advised on initial concept, provided funds for data collection, and gave comments on all drafts of this paper.

## Acknowledgements

The authors would like to thank Robert Bagchi for helpful comments on the manuscript. This work was funded by two Rea Fellowships awarded by the Carnegie Museum of Natural History to K.E.B. Funding for field assistance was provided by a Research Growth Initiative from the University of Wisconsin-Milwaukee to S.A.S. Additional funding for K.E.B. was provided by the Ivy Balsam-Milwaukee Audobon Society Grant. The authors would also like to thank Arie Hunt and Joe Strini for field assistance, Jacob Slyder and James Whitacre for GIS and GPS assistance, Cokie Lindsay for administrative and tactical support, and John Wenzel for input on study design and statistical efforts as well as general support. Thanks also to M. Elizabeth Rodriguez Ronderos, Sergio Estrada Villegas, and Sasha Wright for comments on early drafts of this manuscript.

---

**Table S1:**
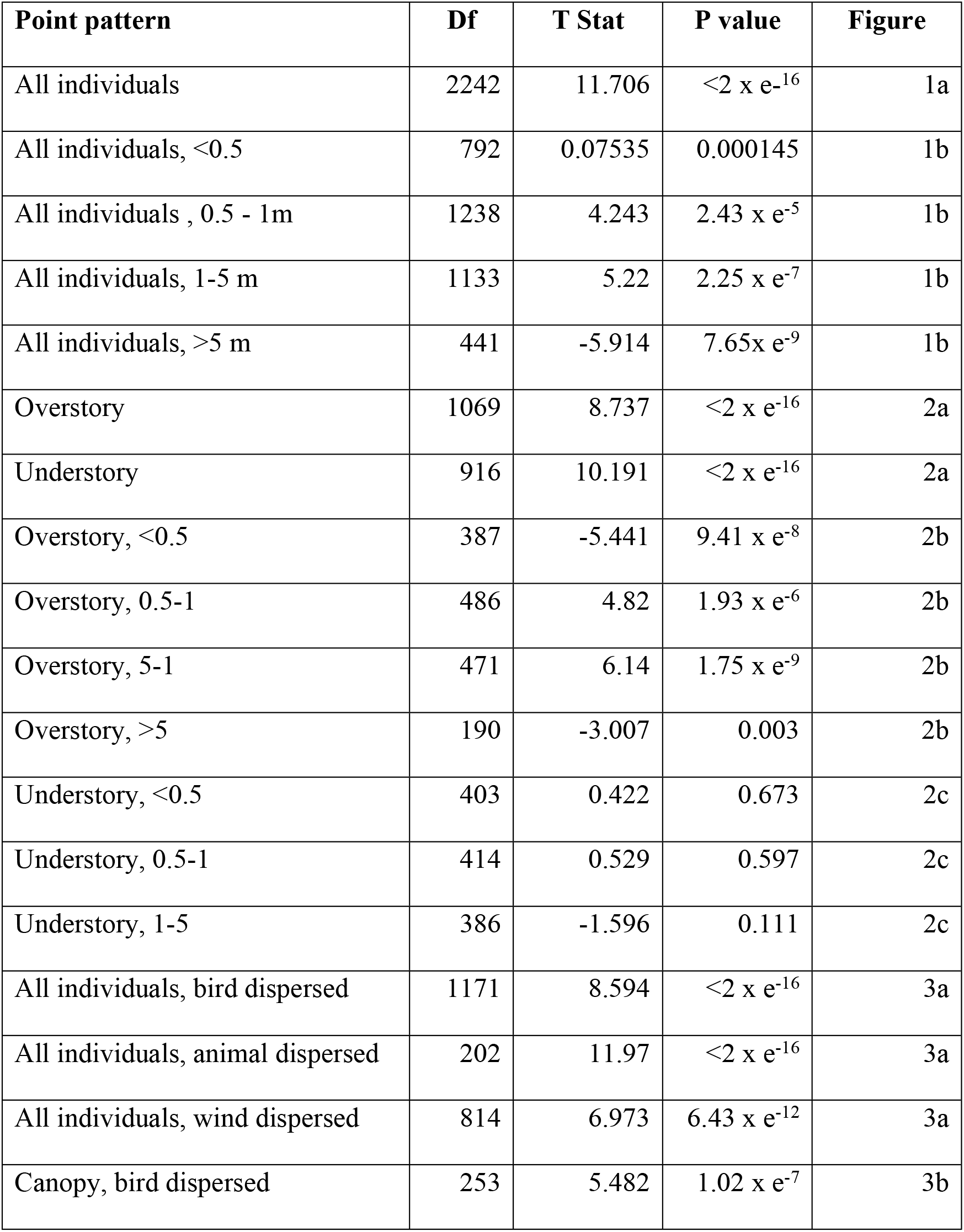
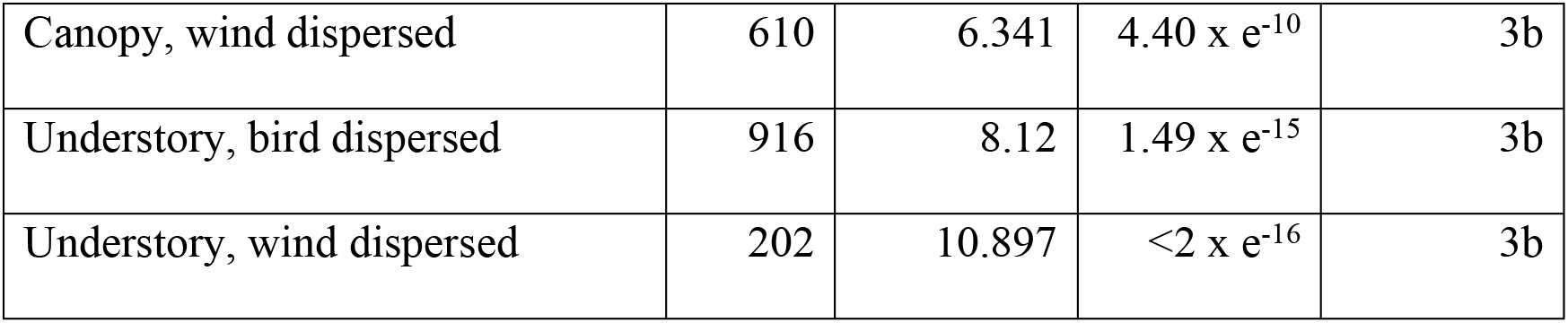
**Linear estimates of the relationship between L and distance for all pooled point patterns at Powdermill Nature Reserve.** We report any pooled point pattern as overdispersed if it has a significantly positive slope and any pooled point pattern as clustered if it has a significantly negative slope.

| Point pattern | Df | T Stat | P value | Figure |
| --- | --- | --- | --- | --- |
| All individuals | 2242 | 11.706 | $<2 \times e^{-16}$ | 1a |
| All individuals, $<0.5$ | 792 | 0.07535 | 0.000145 | 1b |
| All individuals , 0.5 - 1m | 1238 | 4.243 | $2.43 \times e^{-5}$ | 1b |
| All individuals, 1-5 m | 1133 | 5.22 | $2.25 \times e^{-7}$ | 1b |
| All individuals, $>5$ m | 441 | -5.914 | $7.65 \times e^{-9}$ | 1b |
| Overstory | 1069 | 8.737 | $<2 \times e^{-16}$ | 2a |
| Understory | 916 | 10.191 | $<2 \times e^{-16}$ | 2a |
| Overstory, $<0.5$ | 387 | -5.441 | $9.41 \times e^{-8}$ | 2b |
| Overstory, 0.5-1 | 486 | 4.82 | $1.93 \times e^{-6}$ | 2b |
| Overstory, 5-1 | 471 | 6.14 | $1.75 \times e^{-9}$ | 2b |
| Overstory, $>5$ | 190 | -3.007 | 0.003 | 2b |
| Understory, $<0.5$ | 403 | 0.422 | 0.673 | 2c |
| Understory, 0.5-1 | 414 | 0.529 | 0.597 | 2c |
| Understory, 1-5 | 386 | -1.596 | 0.111 | 2c |
| All individuals, bird dispersed | 1171 | 8.594 | $<2 \times e^{-16}$ | 3a |
| All individuals, animal dispersed | 202 | 11.97 | $<2 \times e^{-16}$ | 3a |
| All individuals, wind dispersed | 814 | 6.973 | $6.43 \times e^{-12}$ | 3a |
| Canopy, bird dispersed | 253 | 5.482 | $1.02 \times e^{-7}$ | 3b |
| Canopy, wind dispersed | 610 | 6.341 | $4.40 \times e^{-10}$ | 3b |
| Understory, bird dispersed | 916 | 8.12 | $1.49 \times e^{-15}$ | 3b |
| Understory, wind dispersed | 202 | 10.897 | $<2 \times e^{-16}$ | 3b |

**Table S2:**
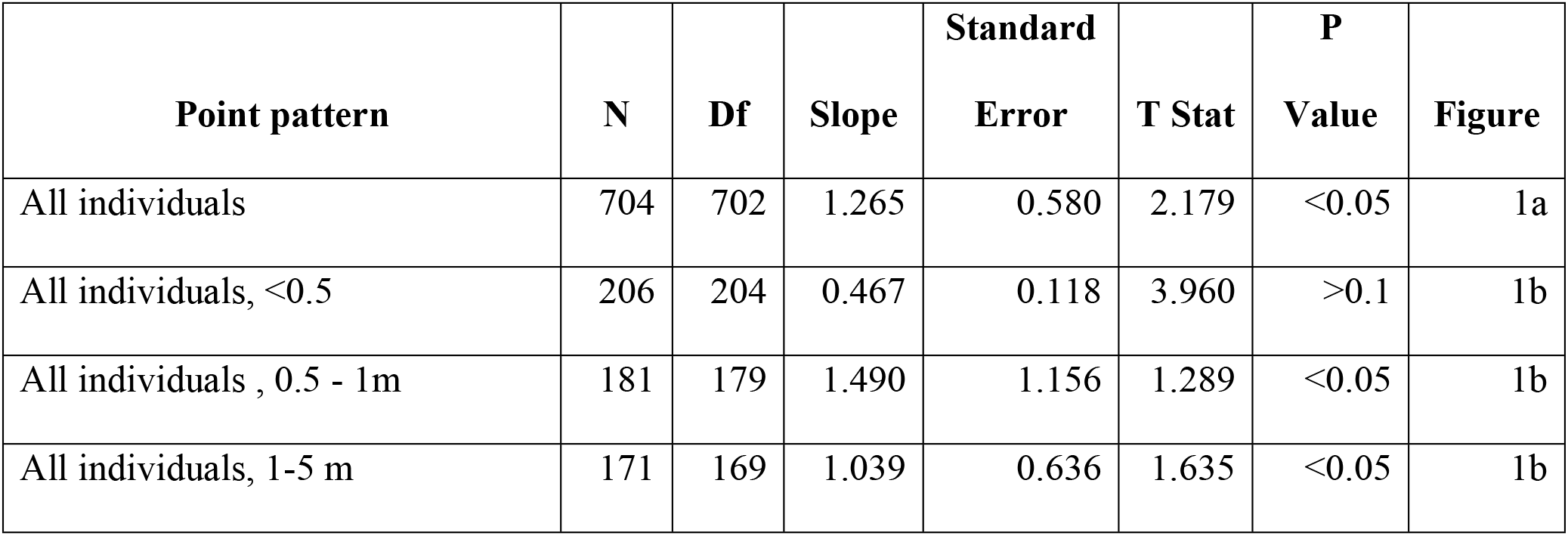
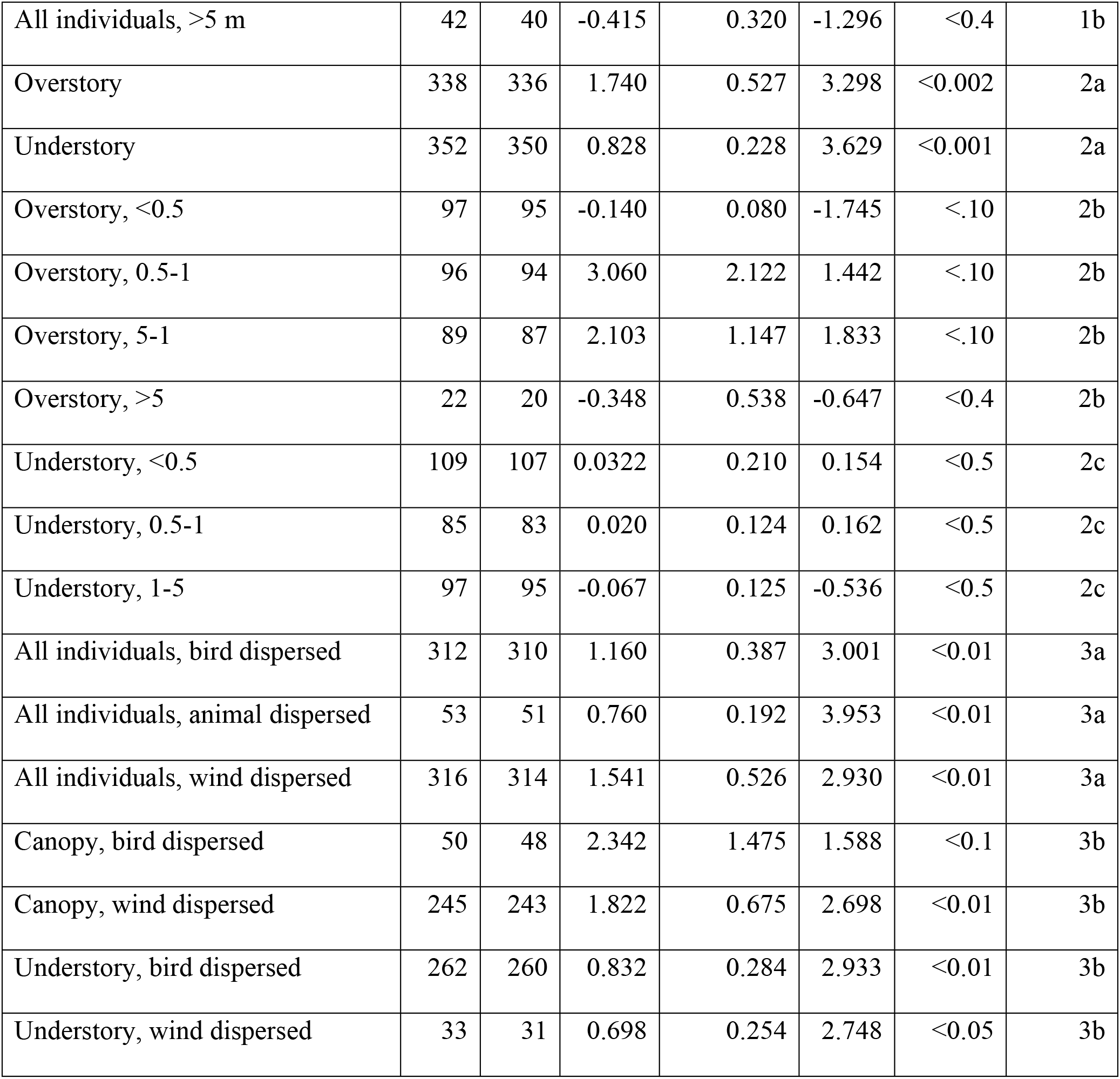
**Linear estimates of the relationship between L and distance for all pooled point patterns utilizing degrees of freedom based on the number of points represented by each point pattern rather than the degrees of freedom based on the number of L estimates.** To make a more conservative estimate of significance, we calculated the P value for each pooled point pattern using the standard deviation of the L estimates and the number of points contributing to each point pattern. We used the number of points contributing rather than the number of L estimates because L is calculated 51 times (each 10 cm distance bin) for each individual point pattern resulting in an inflated degrees of freedom for the overall model. We then calculated the t statistic as the slope/standard error and used a T table to find the estimated P value for a two-tailed t test. We report the P value for each T statistic at the closest degrees of freedom on the table to our degrees of freedom that was not greater than the actual degrees of freedom (i.e. for a degrees of freedom of 204, we report the p-value for 200 degrees of freedom). This analysis may be overly conservative because the variance, standard deviation, and standard error are calculated based on the L estimates which have a higher variance (as they are calculated 51 times per point) than the average L estimate for each point.

**Table S2.**
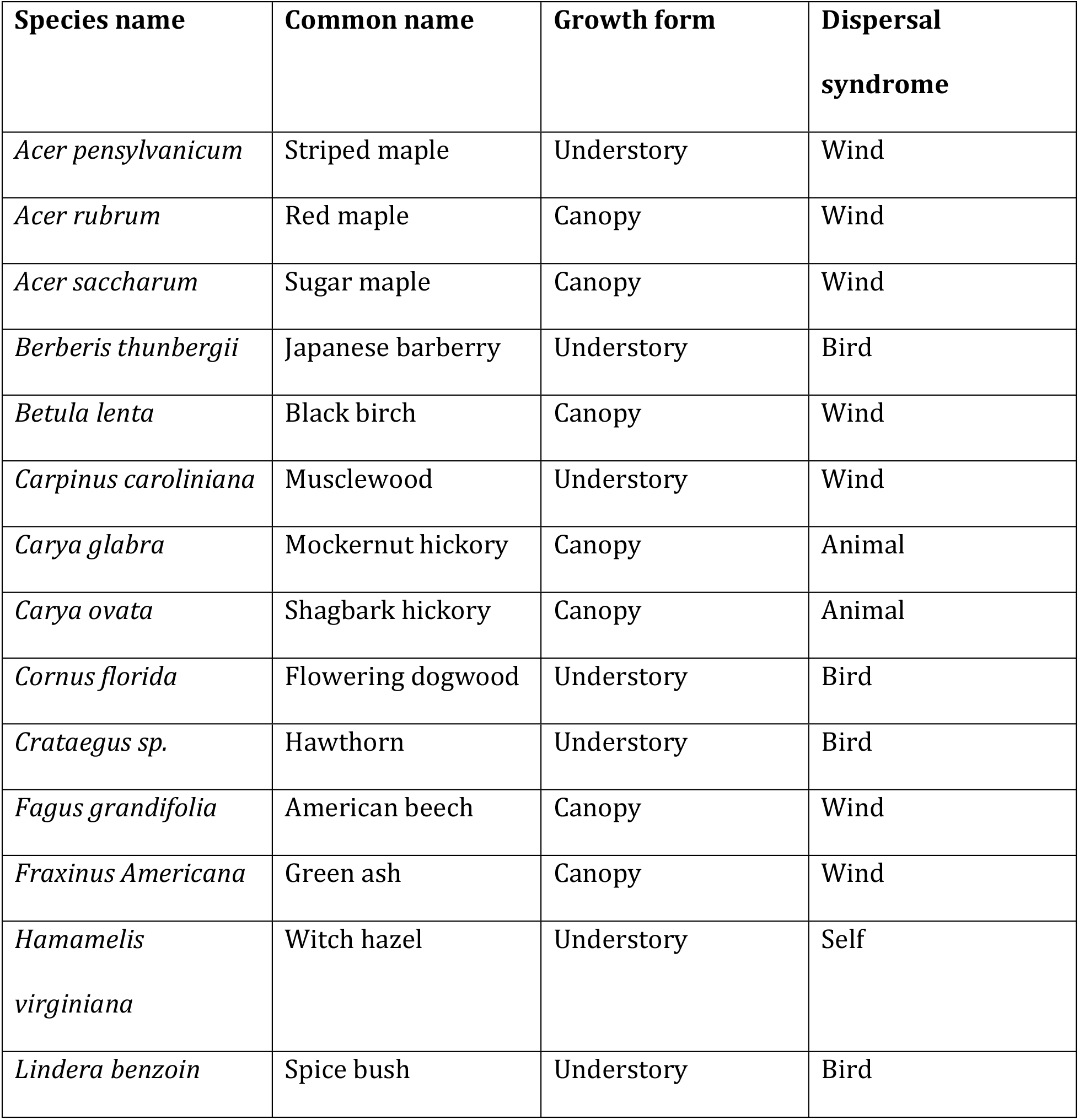
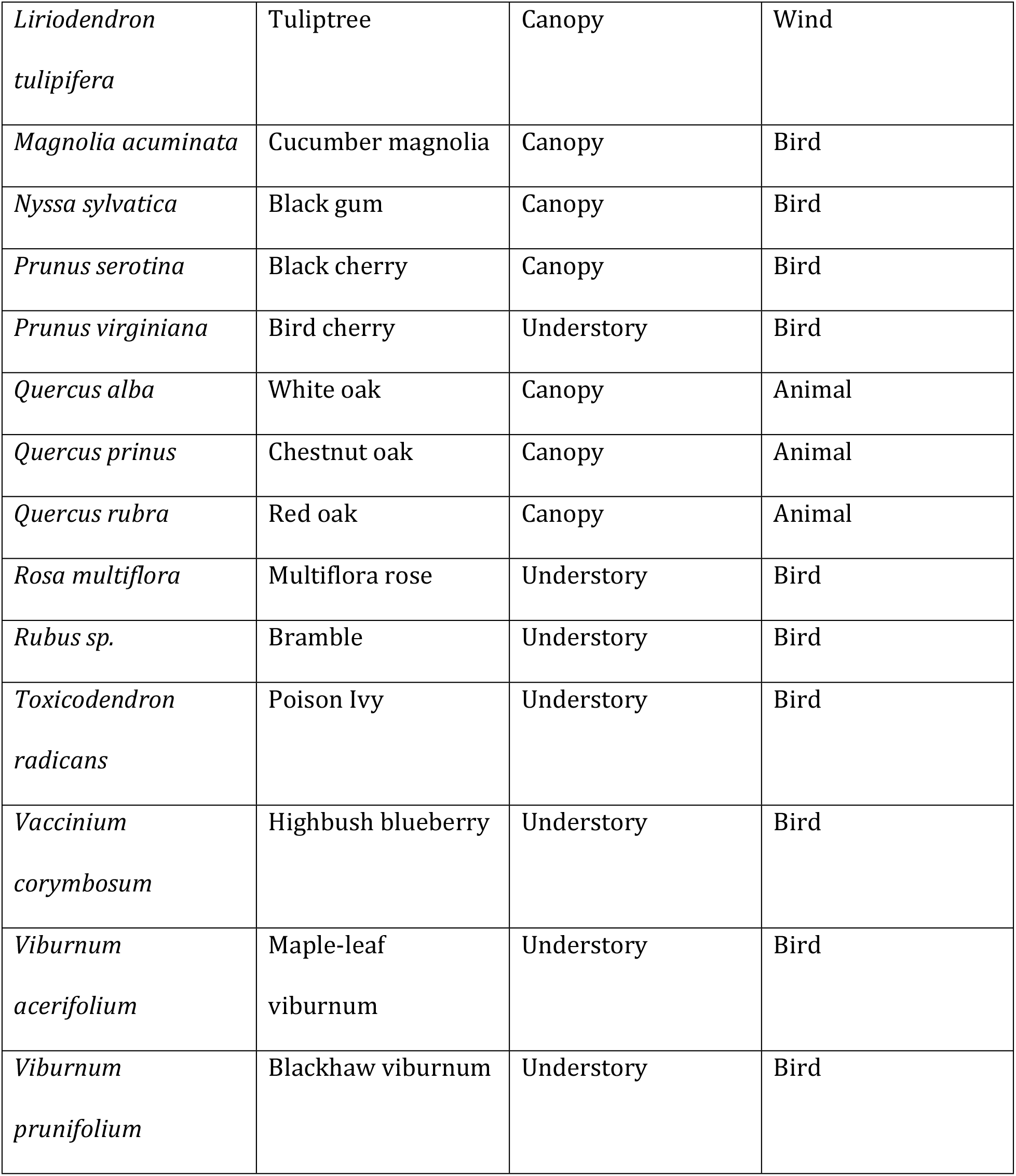
List of species from Powdermill Nature Reserve. We classified species using the Flora of North America species descriptions. If a species had an average height of 5 m or higher, we classified it as a canopy species. If a species had an average height of 5 m or lower, we classified it as an understory species. We based our dispersal syndrome on the description of seed morphology.

